# Biflavonoid quercetin protects against cyclophosphamide–induced organ toxicities via modulation of inflammatory cytokines, brain neurotransmitters, and astrocyte immunoreactivity

**DOI:** 10.1101/2023.01.27.525889

**Authors:** Adejoke Y. Onaolapo, Foluso O Ojo, Olakunle J. Onaolapo

## Abstract

**Background:** Quercetin’s antioxidative properties make it of potential benefit in many clinical conditions where oxidative stress is implicated in the pathological processes.

**Objectives:** To investigate the possible protective effects of quercetin on neurobehaviour, brain oxidative status, markers of inflammation, neurotransmitter balance and astrocyte immunoreactivity in healthy rats administered cyclophosphamide.

**Methods:** Sixty rats were randomly assigned into six groups (n=10). Groups A and D served as normal and cyclophosphamide control respectively and were fed standard rat chow, groups B and E were fed quercetin supplemented diet (100 mg/kg of feed), while those in groups C and F were fed quercetin supplemented diet at 200 mg/kg of feed. Animals in group A-C also received intraperitoneal normal saline on days 1 and 2, while those in groups D-F got intraperitoneal cyclophosphamide (150 mg/kg/day on days 1 and 2). Standard diet and quercetin supplemented diet were administered daily for 21 days. At the end of the experimental period, behavioural tests were carried out, following which animals were sacrificed and blood taken for the assessment of biochemical parameters. Organs were either homogenised or processed for histological study.

**Results:** Quercetin mitigated CYP-induced weight loss and reduction in food consumption. It also improved total antioxidant capacity while reducing CYP-induced lipid peroxidation and biochemical markers of impaired liver function. A reduction in CYP-induced increase in urea and creatinine was also observed. Interleukin-10 increased, while the CYP-induced increase in interleukin-1 beta and TNF-α were also mitigated following quercetin administration. Open field exploratory activities and grooming increased with CYP and the lowest dose of quercetin. The anxiolytic effects of quercetin was demonstrable in the elevated plus maze; while its memory-enhancing effects were seen in the Y-maze and radial-arm maze. Finally, quercetin improved acetylcholine/dopamine levels and brain-derived neurotropic factor, while reducing serotonin levels and astrocyte immunoreactivity.

**Conclusion:** The results show quercetin’s ability to protect against CYP-induced changes in rats. They also highlight the possible use of quercetin as a possible adjunct in cancer chemotherapy-induced tissue damage. However, further research will be needed to delineate its exact role in cancer chemotherapy.

## 1.0 Introduction

Cancer is fast becoming a global major public health challenge, with currently available data showing that it is the second leading cause of death globally (1–3). This is happening despite the fact that currently, early detection and improvement in tools available for management are being shown to help in reducing mortality, at least for some cancers (4). The multimodal approach to cancer management involves the use of chemotherapeutic agents. However, while chemotherapy is crucial to the management of cancers, the severity of the side-effects including cardiotoxicity, neurotoxicity, myelosuppression, hepatotoxicity, and gonadotoxicity may limit use, or result in the discontinuation of therapy (1, 5).

Amongst other factors, an incomplete ability of conventional chemotherapeutic agents to differentiate between normal body cells and cancerous ones is a reason for the side effects. However, this shortcoming (of the drugs) also allows us to further study the effects of such agents in intact biological systems, even in the absence of cancer. In the light of this, it is imperative that research continues towards formulation/identification of adjuvant strategies/agents that are not only effective in preventing or reducing chemotherapy-related side effects, but also do not interfere with the cytotoxic activity of the chemotherapeutic agents (6–9).

Studies (9–15) have continued to support the opinion that a number plants and plant bioactive compounds posses (in addition to their antioxidant and antiinflammatory properties) significant anticancer activity in both *in- vitro* and i*n-vivo* scenarios. Some flavonoids like curcumin and quercetin have also been reported to possess chemopreventive properties through their antioxidant properties; and ability to induce cell-cycle arrest, cell growth and apoptosis (16).There is also recent evidence of the antiproliferative effects of quercetin (either alone or in combination with other flavonoids) in different cancer models (17–20). The main goal of a number of these studies is to provide ample evidence supporting the possible use of plant and plant flavonoids as novel therapies in cancer prevention and treatment. While research continues to gather evidence to support the use of flavonoids like quercetin as anticancer agents, there is an increase in the advocacy for the use of these compounds as adjuvant therapies in cancer management. Such flavonoids or plant compounds could also be beneficial in preventing or mitigating chemotherapy induced toxicities (21). Therefore, this study examined the possible effects of dietary quercetin supplementation on cyclophosphamide induced tissue toxicity in rats. This was with a view to determining the possible benefits of quercetin in mitigating cyclophosphamide induced hepatotoxicity, nephrotoxicity and neurotoxicity in rats.

## 2.0 Materials and Methods

### 2.1 Chemicals and drugs

Quercetin (500 mg, MRM Nutrition, USA), Cyclophosphamide injection (Endoxan-Asta®), Normal saline.

### 2.2 Animals

Healthy male Wistar rats used in this study were obtained from the animal house of the Ladoke Akintola University of Technology Ogbomoso, Oyo State, Nigeria. Rats were housed in wooden cages measuring 20 x 10 x 12 inches in temperature-controlled (22.5°C ±2.5°C) quarters with lights on at 7.00 am. Rats were allowed free access to food and water. All procedures were conducted in accordance with the approved protocols of the Faculty of Basic Medical Sciences, Ladoke Akintola University of Technology and within the provisions for animal care and use prescribed in the scientific procedures on living animals, European Council Directive (EU2010/63).

### 2.3 Diet

All animals were fed commercially available standard rodent chow (29% protein, 11% fat, 58% carbohydrate) from weaning until the commencement of the study. At the beginning of the experimental period, animals were either fed standard chow (29% protein, 11% fat, 58% carbohydrate) or quercetin incorporated into the standard diet at 100 and 200 mg/kg of food and administered *ad libitum* for a period of three weeks.

### 2.4 Experimental methodology

Sixty young adult male rats weighing 120—150 g each were randomly assigned into six groups of ten (n=10) animals each. Group A and D served as normal control and cyclophosphamide control respectively and were fed standard rat chow, rats in groups B, and E were fed quercetin supplemented diet at 100 mg/kg of feed, while those in groups C and F were fed quercetin supplemented diet at 200 mg/kg of feed. Animals in group A-C also received intraperitoneal injection of normal saline at 2ml/kg on days 1 and 2, while those in groups D-F were administered intraperitoneal injection of cyclophosphamide at 150 mg/kg/day on days 1 and 2. Standard diet and quercetin supplemented diet were administered daily for 21 days. At the end of the experimental period, animals were exposed to the behavioural paradigm (open field, Y-maze and elevated plus maze). 24 hours after the behavioural tests animals were sacrificed by cervical dislocation and blood taken through an intracardiac puncture for the assessment of alanine transaminase, aspartate transaminase activity;urea and creatinine levels;as well as levels of inflammatory markers, lipid peroxidation and antioxidant status. The liver, kidneys and brain were removed, observed grossly, weighed and either homogenised or fixed in 10 % neutral buffered formalin. Sections of the liver, kidneys and cerebral cortex were processed for paraffin-embedding, cut at 5 μm and stained for histological study. Supernatant from homogenates of the cerebral cortex was used to assess the activity of brain neurotransmitters

### 2.5 Determination of body weight and food intake

Body weight and food intake were assessed weekly and daily respectively using an electronic weighing balance as previously described [22-24]. Relative change in body weight or food intake was calculated for individual animals using the equation shown below, following which results for all animals were computed to find the statistical mean.

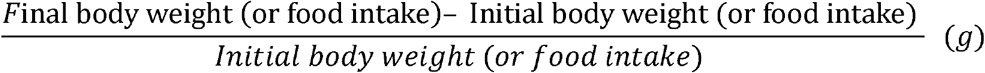

### 2.6 Behavioural tests

Behavioural tests were conducted in the following sequence: 1) elevated plus maze, 2.) open field and 3) memory tests..

#### 2.5.1 Anxiety Model: Elevated plus-maze

The elevated plus-maze which is a plus-shaped apparatus with four arms arranged at right angles to one other was used to measure anxiety-related behaviours. Anxiety behaviours were scored as previously described [25, 26].

#### 2.5.2 Open field behaviours

Open-field responses in rats depict arousal, inhibitory, diversive and inspective exploratory and anxiety behaviours. Also, stereotypic behaviours like grooming have also been represented by this paradigm. These behaviours are generally regarded as central behaviours and are indicative of rodent’s ability to explore. Ten minutes of open field behaviours consisting of grooming, rearing and horizontal locomotion were monitored and scored in the open field box. The open-field paradigm was a rectangular box with a hard floor that measured 36 x 36 x 26 cm. Wood used was painted white, and its floor was divided by permanent red markings into 16 equal squares. Animal placement, movement and scoring system used are as previously described [27, 28].

#### 2.5.3 Memory tests (Y- and Radial arm- maze)

Spatial working memory was assessed using the Y-maze and the radial-arm maze. Spatial working-memory is measured by monitoring spontaneous alternation behaviour. Spontaneous alternation behaviour is the propensity of rodents to alternate conventionally non-reinforced choices of the radial arm maze and/or the Y-maze on successive opportunities. Each rat was placed in one of the arm compartments of the Y maze and allowed to freely move until its tail completely enters another arm. The sequence of arm entries was then recorded as described previously (25, 26, 29).

The radial arm maze apparatus has eight equidistantly-spaced arms that are approximately 3 cm long. Each arm radiates from a small circular central platform.. Each rat is placed on the central platform of the maze and allowed to move freely through respective arms while its behaviours were recorded for 5 min. Working memory was scored when the rat entered each arm a single time as described previously [30, 31]..

### 2.7 Biochemical Test

#### 2.7.1 Estimation of MDA content (Lipid peroxidation)

Lipid peroxidation level was measured as malondialdehyde content as described previously (32–34). Change in colour was measured using a spectrophotometer at 532 nm.

#### 2.7.2 Antioxidant activity

Total antioxidant capacity was determined using commercially available assay kit. Colour changes were measured as described previously described (34–36).

#### 2.7.3 Acetylcholine, dopamine, serotonin and BDNF levels

Supernatants decanted from cerebral cortex homogenate was used to assay for levels of acetylcholine, dopamine, serotonin and brain derived neurotrophic factor using commercially available Enzyme linked immunosorbent assay kits, following the protocols of the manufacturer (ABCAM, Cambridge UK)

#### 2.7.4 Tumour necrosis factor-α, Interleukin (IL) −10 and Interleukin 1β

Tumour necrosis factor-α and interleukin (IL)-10 were measured using enzyme-linked immunosorbent assay (ELISA) techniques with commercially available kits (Enzo Life Sciences Inc. NY, USA) designed to measure the ‘total’ (bound and unbound) amount of the respective cytokines as previously described (37, 38). Interleukin1 β level was assayed using enzyme-linked immunosorbent assay (ELISA) techniques with commercially available kits (Enzo Life Sciences Inc. NY, USA) following the instructions of the manufacturers.

### 2.8 Protocol for Glial fibrillary Acidic Protein (GFAP) Immunohistochemistry

Glial fibrillary acid protein (GFAP) Iimmunohistochemistry was carried out using GFAP primary monoclonal antibody and the Novocastra™ and Novolink DM polymer detection system (Leica Biosystems, UK) as described previously (39–42).

### 2.9 Photomicrography

Histologically processed sections of the liver, kidneys and cerebral cortex were examined microscopically using a Sellon-Olympus trinocular microscope (XSZ-107E, China) with a digital camera (Canon Powershot 2500), and photomicrographs taken. Histopathological changes were assessed by a pathologist that was blinded to the groupings.

### 2.10 Statistical analysis

Data were analysed with Chris Rorden’s ANOVA for windows (version 0.98). Data analysis was by One-way analysis of variance (ANOVA) and post-hoc test (Tukey HSD) was used for within and between group comparisons. Results were expressed as mean ± S.E.M. and p < 0.05 was taken as the accepted level of significant difference from control.

## 3.0 Result

### 3.1 Effect of quercetin on body weight

Figure 1 shows the effect of dietary quercetin supplementation on weekly body weight (upper panel) and percentage change in body weight (lower panel) in rats exposed to cyclophosphamide (CYP). There was a significant (p <0.001) decrease in weekly weight gain in the groups administered CYP, CYP+ quercetin (CYP/Q) at 100 and 200 mg/kg of feed compared to control. Compared to CYP, body weight increased with CYP/Q100 and CYP/Q200. Over the treatment period, percentage weight gain was observed to decrease in the CYP groups *CYP (−44.4%). CYP+Q100 (14.3%) and CYP+ Q 200 (−4.4%) compared to control (26.2%). Compared to CYP (−44.4%), weight gain was better with CYP+Q100 (−14.3%) and CYP+ Q 200 (−4.4%) respectively.

**Figure 1:**
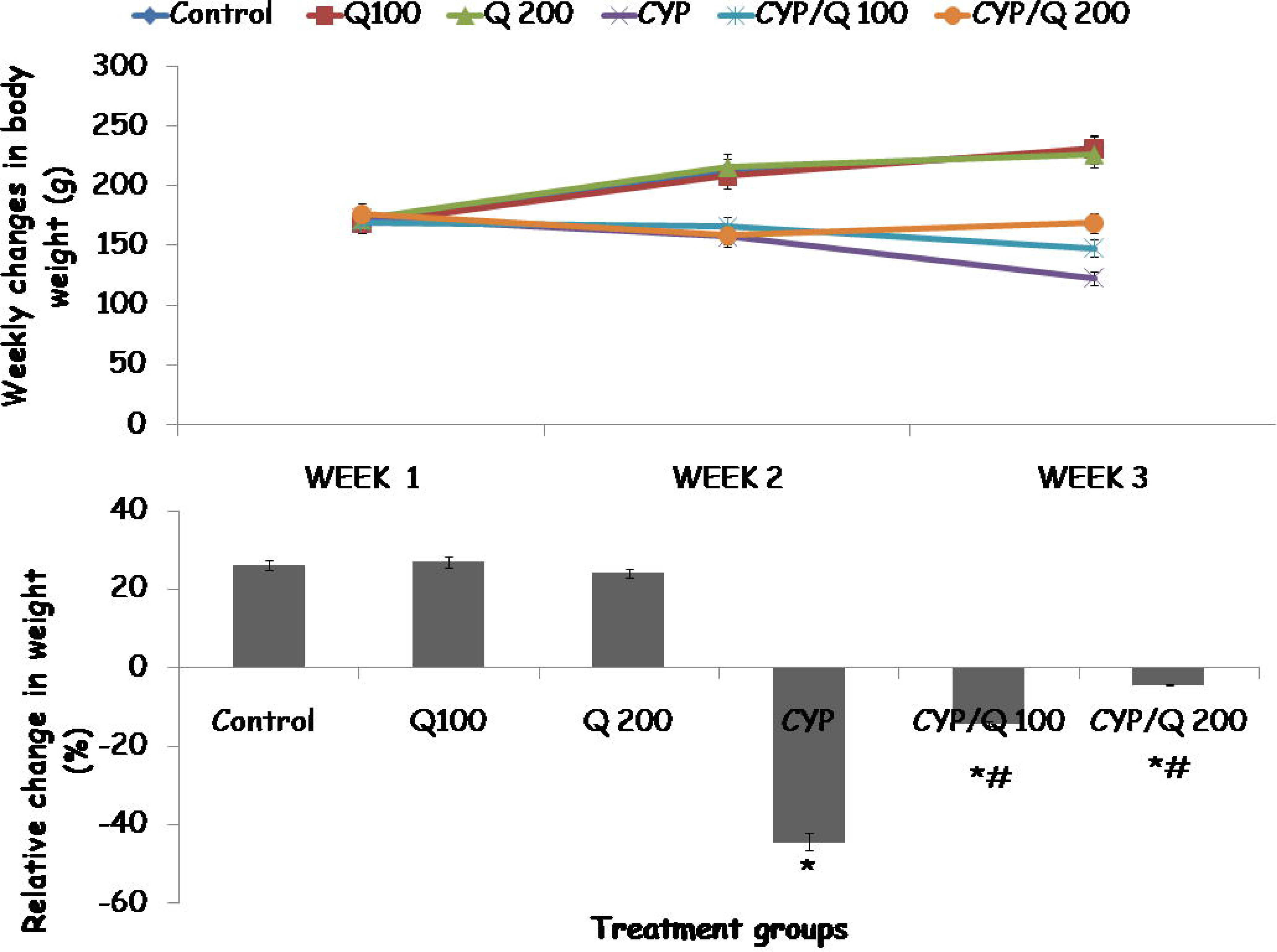
Effect of quercetin (Q) on weekly body weight (upper panel) and relative change in weight (lower panel) in CYP treated rats. Each bar represents Mean ± S.E.M, *p < 0.05 vs. control, # p<0.05 significant difference from CYP, number of mice per treatment group =10. CYP: Cyclophosphamide, Q: Quercetin.

### 3.2 Effect of quercetin on food intake

Figure 2 shows the effect of dietary quercetin supplementation on weekly food intake and percentage weight gain (lower panel) in rats exposed to CYP. There was a significant (p < 0.001) decrease in food intake with CYP, CYP/Q 100 and 200 compared to control. Compared to CYP, weekly food intake increased with CYP/Q100 and CYP/Q200 respectively. Over the treatment period, percentage change in food intake decreased with CYP (−90.8%). CYP+Q100 (−16.8%) and CYP+ Q 200 (−8.01%) compared to control (51.5%). Compared to CYP (−90.8%), food intake increased with CYP+Q100 (−16.0%) and CYP+ Q 200 (−8.01%) respectively.

**Figure 2:**
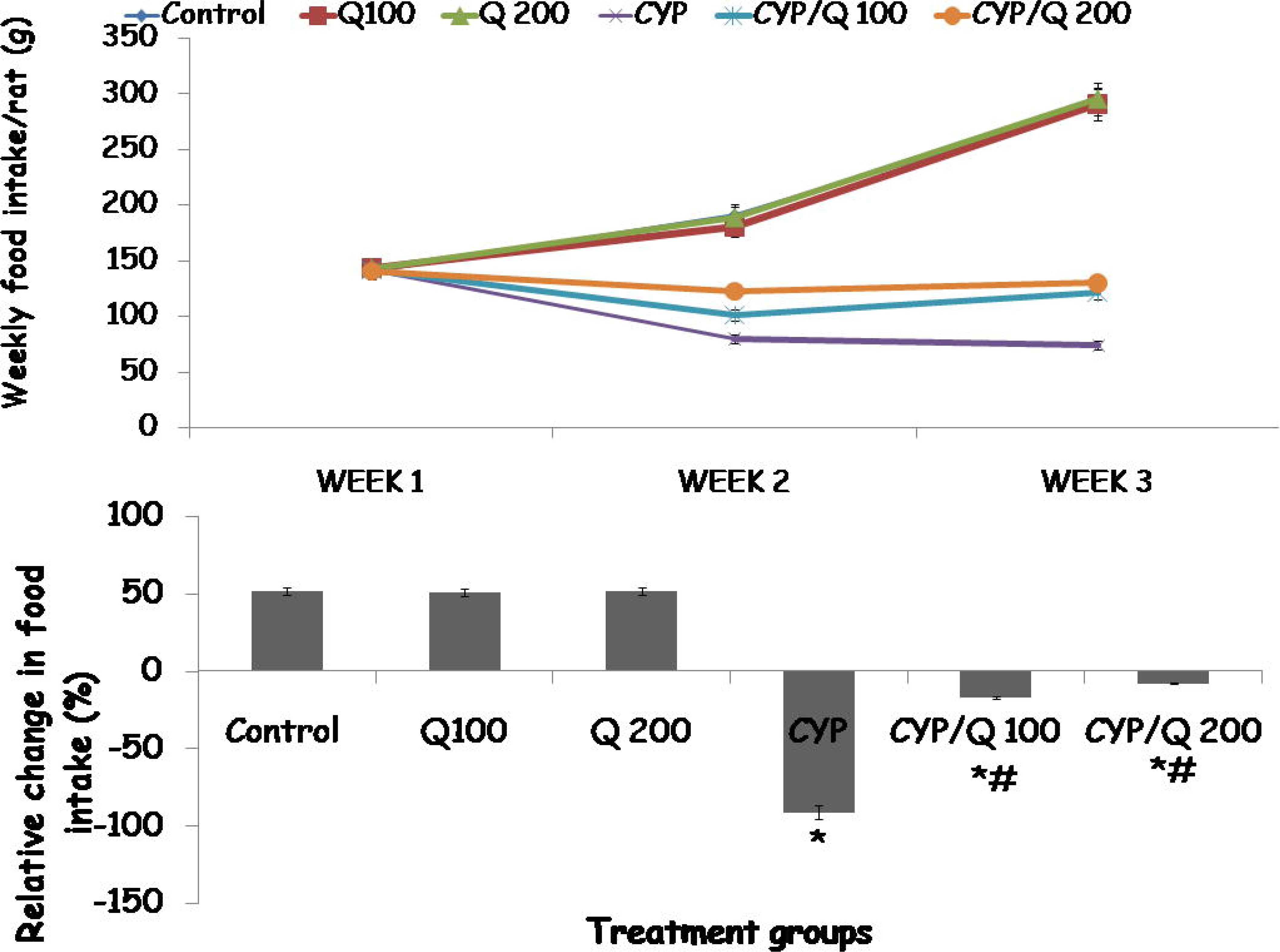
Effect of quercetin (Q) on weekly food intake (upper panel and percentage increase in food intake (lower panel) in rats exposed to CYP. Each bar represents Mean ± S.E.M, *p < 0.05 vs. control, # p<0.05 significant difference from CYP, number of mice per treatment group =10. CYP: Cyclophosphamide, Q: Quercetin.

### 3.3 Effect of dietary quercetin supplementation on liver transaminase activity and oxidative stress parameters

Table 1 shows effect of dietary quercetin supplementation on liver transaminase activity, total antioxidant capacity and malondialdehyde levels in CYP treated rats. Activities of aspartate (AST) and alanine transaminase (ALT) increased significantly with CYP, CYP/Q100, and CYP/Q 200 compared to control respectively. Compared to CYP, activities of AST and ALT increased with CYP/Q100 and CYP/Q200.

**Table 1:** Effect of quercetin on liver transaminase levels, lipid peroxidation, and antioxidant capacity in CYP-treated rats.

| Groups | AST (IU/L) | ALT (IU/L) | MDA uM | TAC (TE mM) |
| --- | --- | --- | --- | --- |
| Control | 80.31±0.14 | 40.01 ±1.13 | 6.05±0.10 | 14.15±0.10 |
| Q100 | 82.41±0.22 | 42.09 ±1.22 | 5.95±0.12 | 25.65±0.12* |
| Q200 | 83.10±0.33 | 42.11 ±1.10 | 5.82±0.14 | 27.85±0.12* |
| CYP | 113.20±0.40* | 88.10±1.29* | 22.60±0.30 | 8.22±0.15 |
| CYP/Q100 | 99.40±0.22* | 66.20±1.22* | 18.58±0.30*# | 16.30±0.10* |
| CYPQ200 | 87.20±0.62* | 60.40±1.31* | 15.35±0.22*# | 15.23±0.20* |

Data presented as Mean ± S.E.M, \*p < 0.05 vs. control, #p<0.05 significant difference from CYP, number of rats/group=10, AST: Aspartate transaminase, ALT: Alanine transaminase, MDA: malondialdehyde, TAC: Total antioxidant capacity.

Total antioxidant capacity (TAC) increased significantly with Q100 and Q200 while it decreased significantly with CYP, CYP/Q100 and CYP/Q200 compared to control respectively. Compared to CYP, TAC increased with CYP/Q100 and CYP/Q200.

Malondialdehyde (MDA) levels increased significantly with CYP, CYP/Q100 and CYP Q200 compared to control. Compared to CYP, MDA levels decreased with CYP/Q100 and CYP/Q200.

### 3.4 Effect of dietary quercetin on levels of urea, creatinine, and inflammatory cytokines

Table 2 shows the effect of dietary quercetin supplementation on urea, creatinine, tumour necrosis factor-α, interleukin-10 and interleukin-1β in CYP-treated rats. Level of urea and creatinine increased significantly with CYP, CYP/Q100 and CYP/Q 200 compared to control respectively. Compared to CYP, urea and creatinine levels increased with CYP/Q100 and CYP/Q200. Interleukin-10 levels increased significantly with Q100 and Q200 while it decreased significantly with CYP CYP/Q100 and CYP/Q 200 compared to control. Compared to CYP, interleukin-10 levels increased with CYP/Q100 and CYP/Q200. Interleukin 1β levels increased significantly with CYP, CYP/Q100 and CYP Q200 and decreased with Q100 and Q200 compared to control. Compared to CYP, interleukin 1β levels decreased with CYP/Q100 and CYP/Q200. Tumour necrosis factor α levels increased significantly with CYP and decreased with CYP/Q100 and CYP Q200 compared to control. Compared to CYP, tumour necrosis factor α levels decreased with CYP/Q200.

**Table 2:** Effect of quercetin on levels of urea, creatinine and inflammatory cytokines in CYP-treated rats.

| Groups | Urea<br>(mmol/L) | Creatinine<br>(mmol/L) | Interleukin<br>10 (pg/ml) | Interleukin-<br>1β (pg/ml) | TNF-α(pg/ml) |
| --- | --- | --- | --- | --- | --- |
| Control | 2.69±0.011 | 43.0±0.23 | 59.22±1.20 | 33.04±1.00 | 23.16±0.34 |
| Q100 | 2.42±0.12 | 42.50±0.22 | 60.11±1.32* | 28.25±1.02* | 22.06±0.22 |
| Q200 | 2.56±0.22 | 42.55±0.21 | 65.18±1.52* | 25.12±1.43* | 22.15±0.41 |
| CYP | 31.32±0.14* | 87.50±0.22* | 18.54±1.80* | 82.22±2.20* | 30.20±0.65* |
| CYP/Q100 | 25.96±0.32* | 59.55±0.21*# | 25.54±2.15*# | 48.21±2.20*# | 16.12±0.15*# |
| CYPQ200 | 17.26±0.45* | 50.15±0.24*# | 35.30±2.60*# | 45.24±2.20*# | 15.22±0.30*# |

Data presented as Mean ± S.E.M, \*p < 0.05 vs. control, #p<0.05 significant difference from CYP, number of rats/group =6, TNF-α : Tumour necrosis $\alpha$ .

### 3.5 Effect of quercetin on neurobehavioural indices

#### 3.5.1 Effect of quercetin on open field locomotor activity

Figure 3 shows the effect of dietary quercetin supplementation on open field horizontal locomotion (upper panel) and vertical locomotion (lower panel) in CYP treated rats. Horizontal locomotion increased with Q100, CYP, CYP/Q100 and CYP/Q200 compared to control. Compared to CYP, horizontal locomotion decreased with CYP/Q100 and CYP/Q200.

**Figure 3:**
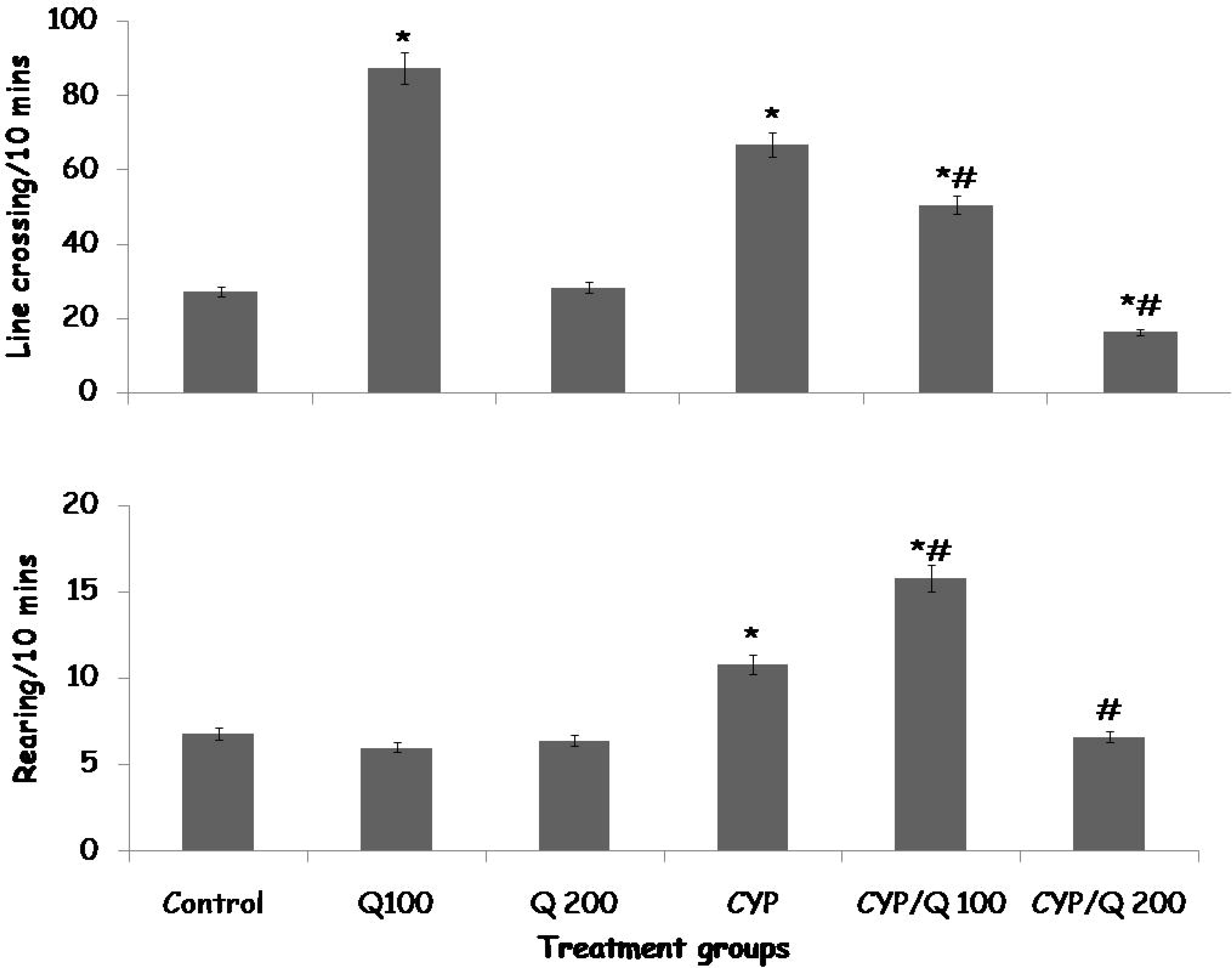
Effect of quercetin (Q) on horizontal locomotion (upper panel) and vertical locomotion (lower panel) in rats treated with CYP. Each bar represents Mean ± S.E.M, *p < 0.05 vs. control, # p<0.05 significant difference from CYP, number of mice per treatment group =10. CYP: Cyclophosphamide, Q: Quercetin.

Vertical locomotion (rearing) increased with CYP and CYP/Q100 compared to control. Compared to groups treated with CYP, rearing increased with CYP/Q100 and decreased with CYP/Q200.

#### 3.5.2 Effect of quercetin on self-grooming and catalepsy test scores

Figure 4 shows the effect of dietary quercetin supplementation on self-grooming (upper panel) and catalepsy bar test scores (lower panel) in CYP treated rats. Self-grooming increased with Q100, CYP and CYP/Q100 and decreased with CYP/Q 200 compared to control. Compared to CYP, there was a significant increase in self-grooming with CYP/Q100, and decreased with CYP/Q200.

**Figure 4:**
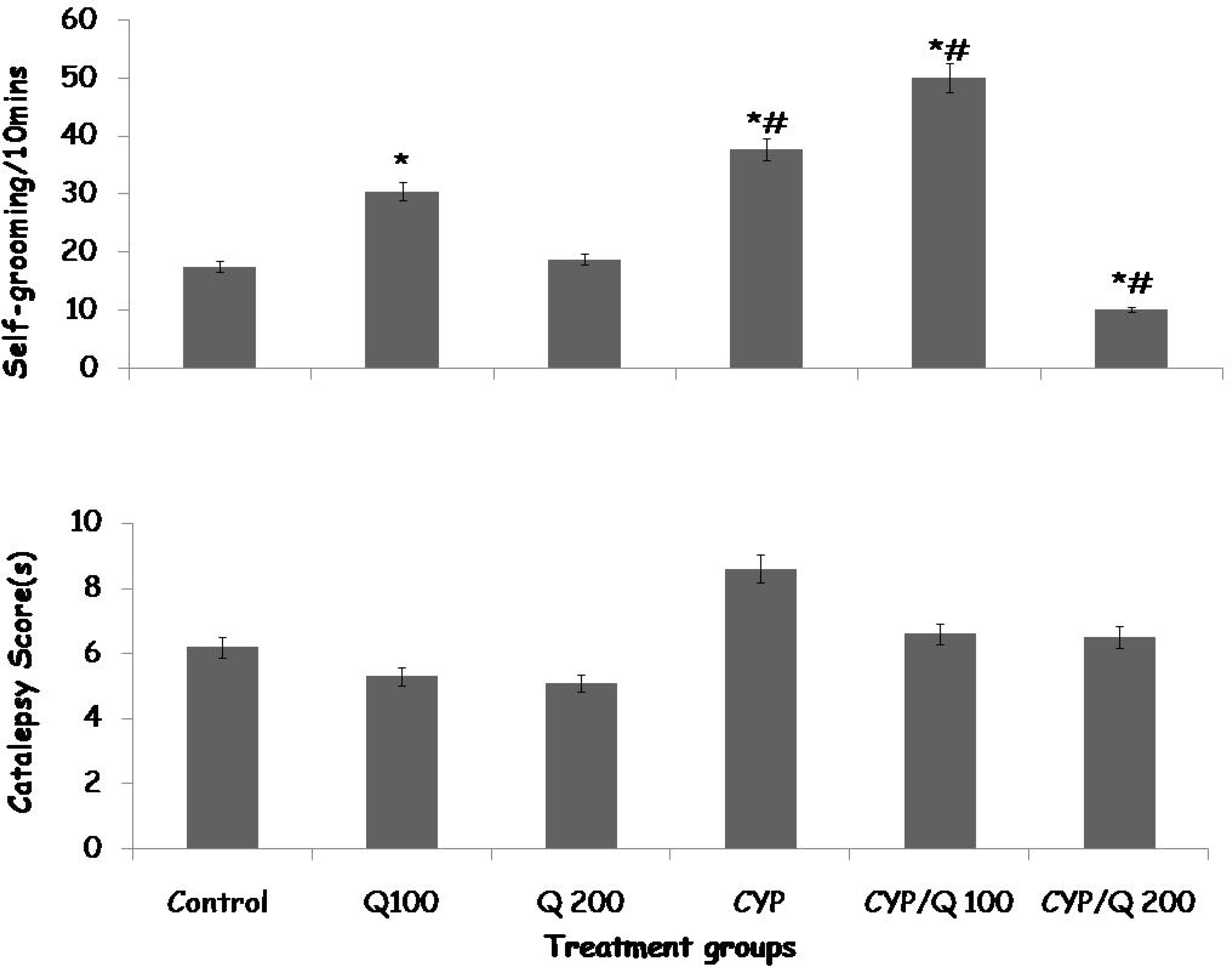
Effect of quercetin (Q) on self −behaviour (upper panel) and catalepsy bar test (lower panel) in rats treated with CYP. Each bar represents Mean ± S.E.M, *p < 0.05 vs. control, # p<0.05 significant difference from CYP, number of mice per treatment group =10. CYP: Cyclophosphamide, Q: Quercetin.

**Figure 5:**
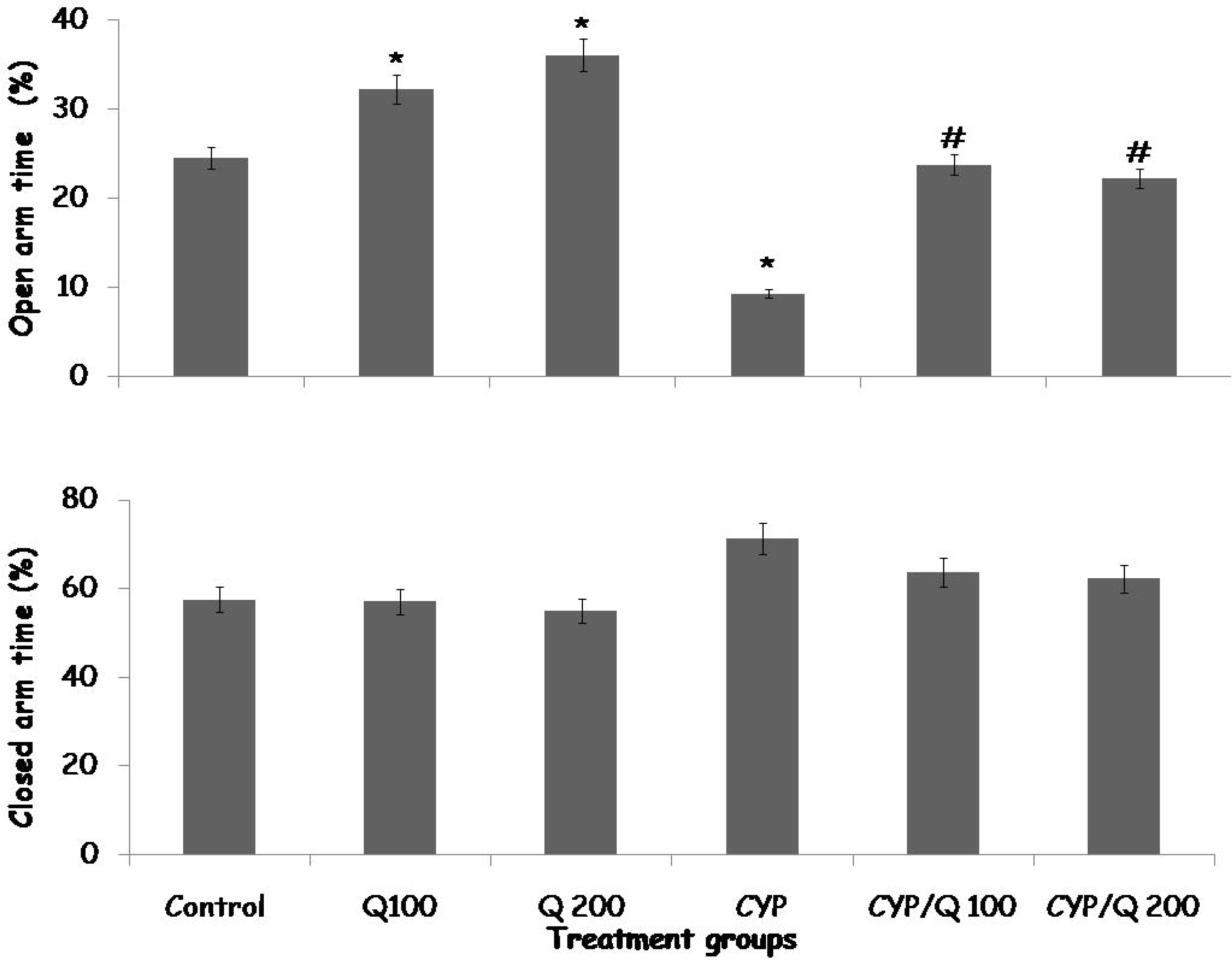
Effect of quercetin (Q) on percentage open arm time (upper panel) and percentage closed arm time (lower panel) in rats treated with CYP. Each bar represents Mean ± S.E.M, *p < 0.05 vs. control, # p<0.05 significant difference from CYP, number of mice per treatment group =10. CYP: Cyclophosphamide, Q: Quercetin.

Catalepsy bar test scores did not differ significantly in any of the groups administered quercetin and/or CYP compared to control or CYP.

#### 3.5.3 Effect of quercetin on open and closed arm times in the elevated plus maze

Figure 4 shows the effect of dietary quercetin supplementation on percentage open arm time (upper panel) and closed arm time (lower panel) in CYP treated rats. Open arm time increased with Q100 and Q200 and decreased with CYP compared to control. Compared to CYP, open arm time increased with CYP/Q100 and CYP/Q200.

Elevated plus maze closed arm time did not differ significantly in any of the groups administered quercetin and/or CYP compared to control or CYP.

#### 3.5.4 Effect of quercetin on Spatial working memory in the Y and Radial arm maze

Figure 6 shows the effect of dietary quercetin supplementation on Y maze (upper panel) and radial arm maze (lower panel) spatial working memory in CYP treated rats. Percentage alternation in the Y maze increased with Q100, Q200, CYP/Q100 and CYP/Q 200 and decreased with CYP compared to control. Compared to CYP, Y maze alternation increased with CYP/Q100 and CYP/Q200.

Alternation index in the radial arm maze increased with Q100, CYP/Q100 and CYP/Q200 compared to control. Compared to groups treated with CYP, radial arm maze alternation index increased with CYP/Q100 and CYP/Q200.

**Figure 6:**
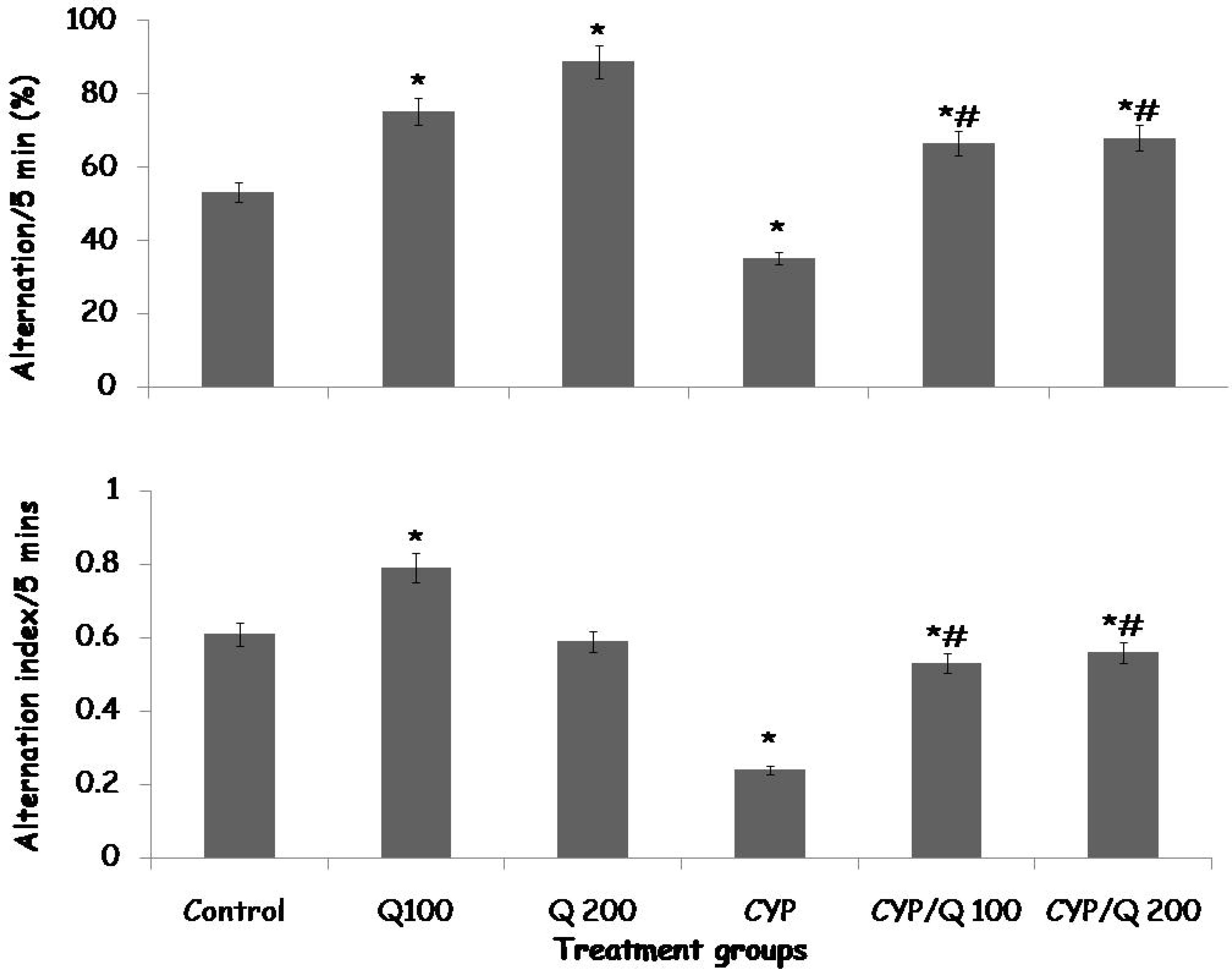
Effect of quercetin (Q) on Y maze (upper panel) and radial arm maze (lower panel) spatial working memory in rats treated with CYP. Each bar represents Mean ± S.E.M, *p < 0.05 vs. control, # p<0.05 significant difference from CYP, number of mice per treatment group =10. CYP: Cyclophosphamide, Q: Quercetin.

### 3.6 Effect of quercetin on acetylcholine, dopamine, serotonin and brain derived neurotropic factor levels

Table 3 shows the effect of dietary quercetin supplementation on neurotransmitter levels in the cerebral cortex in CYP treated rats. There was a significant increase in acetylcholine levels with Q100 and Q200 and a decrease with CYP, CYP/Q100 and CYP/Q 200 compared to control. Compared to CYP, acetylcholine levels increased with CYP/Q100 and CYP/Q200.

**Table 3:** Effect of quercetin on cerebral cortex neurotransmitter levels.

| Groups | Acetylcholine<br>(pmol/mg tissue) | Dopamine<br>(pmol/mg tissue) | BDNF<br>(pmol/mg<br>tissue) | Serotonin<br>(pmol/mg<br>tissue) |
| --- | --- | --- | --- | --- |
| Control | 3.50±0.01 | 45.10±1.11 | 23.01±0.10 | 8.53±0.20 |
| Q100 | 6.12±0.09* | 47.12±1.02 | 32.11±0.12* | 8.21±0.10* |
| Q200 | 7.22±0.10* | 49.34±1.01 | 35.10±0.11* | 8.62±0.33* |
| CYP | 0.82±0.02* | 22.12±0.78* | 12.50±0.80* | 17.10±1.34* |
| CYP/Q100 | 1.22±0.02* | 32.77±0.89*# | 15.25±0.10*# | 10.32±2.16# |
| CYPQ200 | 2.01±0.01* | 41.00±0.20*# | 15.23±0.65*# | 9.43±1.14# |

Data presented as Mean ± S.E.M, \*p < 0.05 vs. control, #p<0.05 significant difference from CYP, number of rats/group =6, BDNF: Brain derived neurotropic factor.

Dopamine levels increased significantly with Q100 and Q200 and decreased with CYP, CYP/Q100 and CYP/Q200 compared to control. Compared to CYP, dopamine levels increased with CYP/Q100 and CYP/Q200.

Brain derived neurotropic factor (BDNF) levels increased significantly with with Q100 and Q200 and a decrease with CYP, CYP/Q100 and CYP/Q200 compared to control. Compared to CYP, BDNF levels increased with CYP/Q100 and CYP/Q200.

Serotonin levels were significantly increased with CYP, compared to control. Compared to CYP, serotonin levels increased with CYP/Q100 and CYP/Q200.

### 3.7 Effect of quercetin on kidney and liver histomorphology

Examination of haematoxylin and eosin stained sections of the kidney of rats in the control group (Figure 7a) and in groups fed quercetin diet (Figure 7b, 7c) revealed well-demarcated cortex and medulla. The Bowman’s capsule, glomeruli, proximal and distal renal tubules and blood vessels appeared within normal limits. The glomerular and tubular nuclei were also deeply staining. Examination of sections of the kidney obtained from animals in the CYP group (Figure 7d) revealed a disruption of normal kidney architecture with widening of the Bowman’s space, contraction of the renal glomeruli and dilatation of the proximal and n distal renal tubules. Pale-staining tubular and glomerular nuclei were also observed. In groups treated with quercetin (Figure 7e, 7f), protection from kidney injury was observed.

**Figure 7 (a–f):**
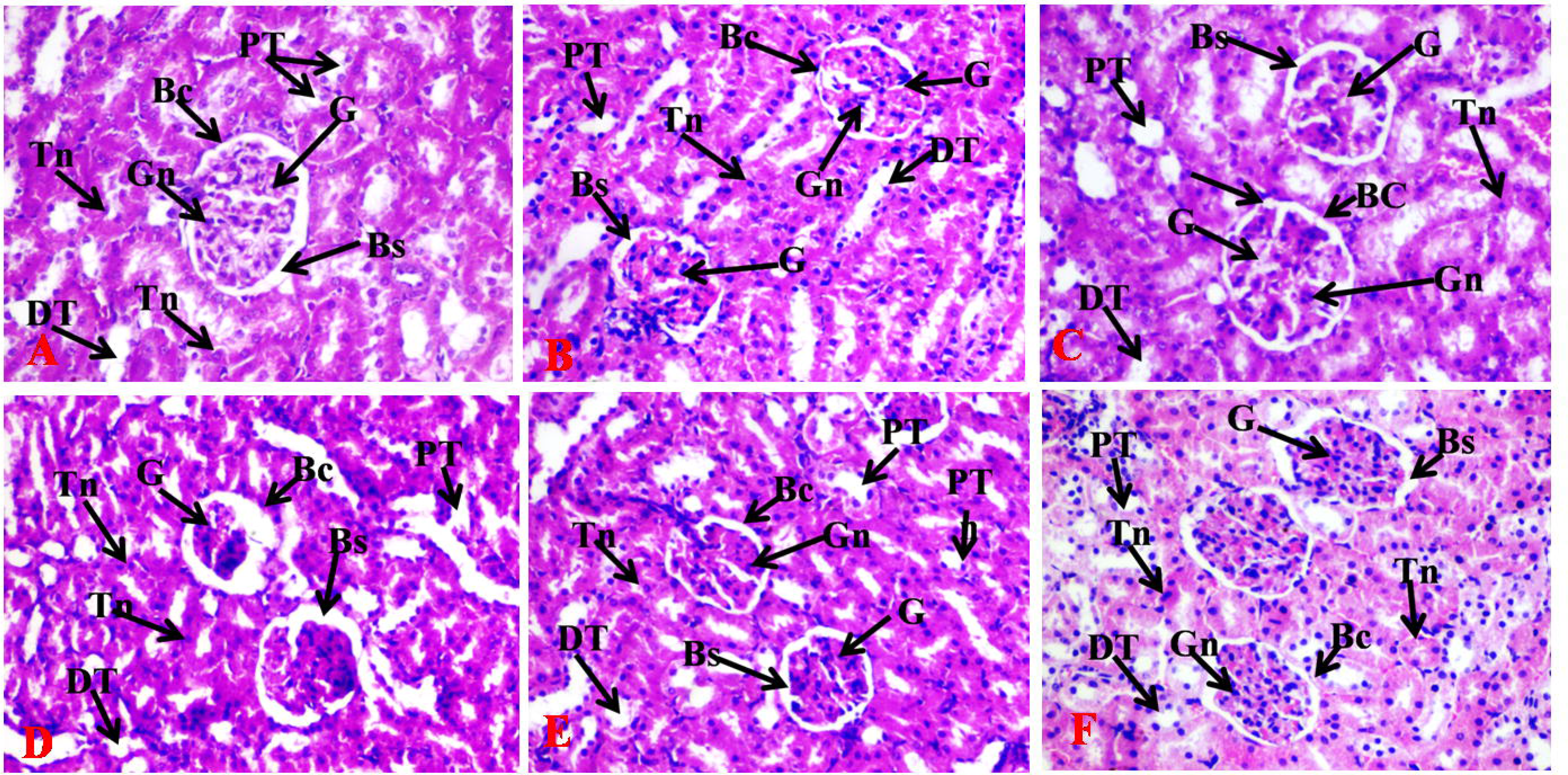
Effect of quercetin on kidney morphology in cyclophosphamide (CYP)-administered rats. 7a) Control, 7b) Quercetin at 100 mg/kg body weight, 7c) Quercetin at 200 mg/kg body weight, 7d) Cyclophosphamide (CYP), 7e) CYP plus Quercetin at 100 mg/kg body weight, 7f) CYP plus Quercetin at 200 mg/kg body weight. Representative photomicrographs of haematoxylin and eosin (H&E) stained sections of the rat kidney show the glomerulus (G), Bowman’s capsule (BC) Bowman’s space (BS), proximal tubule (PT), and distal tubule (DT), tubular nuclei (Tn) and glomerular nuclei (Gn). H&E 160

Examination of haematoxylin and eosin-stained sections of the liver of rats in the control group (Figure 8a) and those fed quercetin (figure 8b, 8c) revealed sheets of radially-arranged hepatocytes around the central vein, with hepatic sinusoids demarcating cords of hepatocytes. Hepatocyte nuclei were deeply staining; these features were in keeping with normal liver histology. In the CYP group (Figure 8d) marked disruption of hepatocyte architecture, with loss of intervening sinusoidal spaces, swollen hepatocytes and pale-staining hepatocyte nuclei were observed; these features are in keeping with hepatic injury. In groups treated with quercetin (Figure 8e & 8f) a protection against CYP induced hepatotoxicity was observed.

**Figure 8(a–f):**
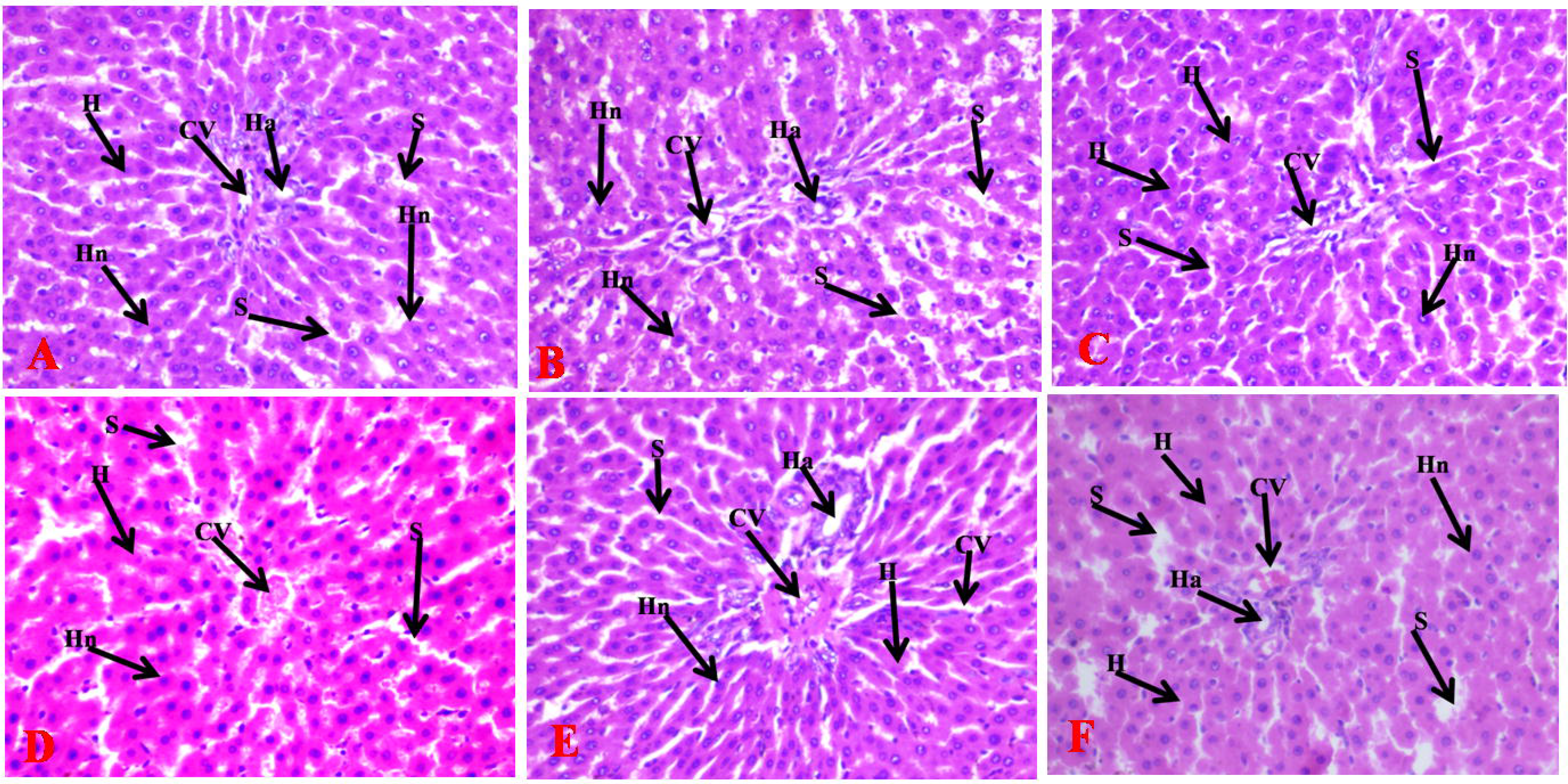
Effect of quercetin on liver morphology in Cyclophosphamide-administered rats 8a) Control, 8b) Quercetin at 100 mg/kg body weight, 8c) Quercetin at 200 mg/kg body weight, 8d) Cyclophosphamide (CYP), 8e) CYP plus Quercetin at 100 mg/kg body weight, 8f) CYP plus Quercetin at 200 mg/kg body weight. Representative photomicrographs of haematoxylin and eosin (H&E) stained sections of the rat liver show sinusoidal spaces (S) demarcating hepatocytes (H), with hepatocyte nucleus (Hn), Central vein (CV), hepatic artery (Ha) are also observed. H&E 160.

### 3.8 Cerebral cortex histomorphology

Figures show representative haematoxylin and eosin (figure 9a-f) and cresyl fast violet (figure 10a-f) stained sections of the rat cerebral cortex. Examination of the haematoxylin and eosin stained slides revealed distinct layers of the cerebral cortex with presence of numerous granule cells, pyramidal cells and neuroglia in rats administered vehicle (Fig. 9a-c). Multipolar shaped pyramidal cells with large, rounded, vesicular nucleus are seen scattered throughout the neuropil; granular neurons with large open-faced nuclei, prominent nucleoli and scant cytoplasm are also observed. The pink-staining background which is the neuropil is better appreciated in the cresyl fast violet stained slides (figure 10 a-c). In the group administered CYP, numerous degenerating pyramidal and granule cells are observed, degenerating neurons are pale-staining with shrunken pale-staining nuclei, these features are in keeping with neuronal injury. In groups treated with quercetin (9d-e) was observed a protection against CYP-induced neuronal injury. Examination of cresyl-fast violet stained sections of the cerebrum revealed characteristic layered arrangement of the cerebrum with well-delineated multipolar pyramidal cells, deeply-staining granule cells and neuroglia in the control group (figure 10a) and groups administered quercetin (Fig. 10b, 9c). However, degenerating pyramidal cells with pale staining nuclei were observed in the group administered CYP (Figure 10d), while a protection against neuronal injury was observed in the groups treated with quercetin (Figure 10 e, 10 f).

**Figure 9 (a–f):**
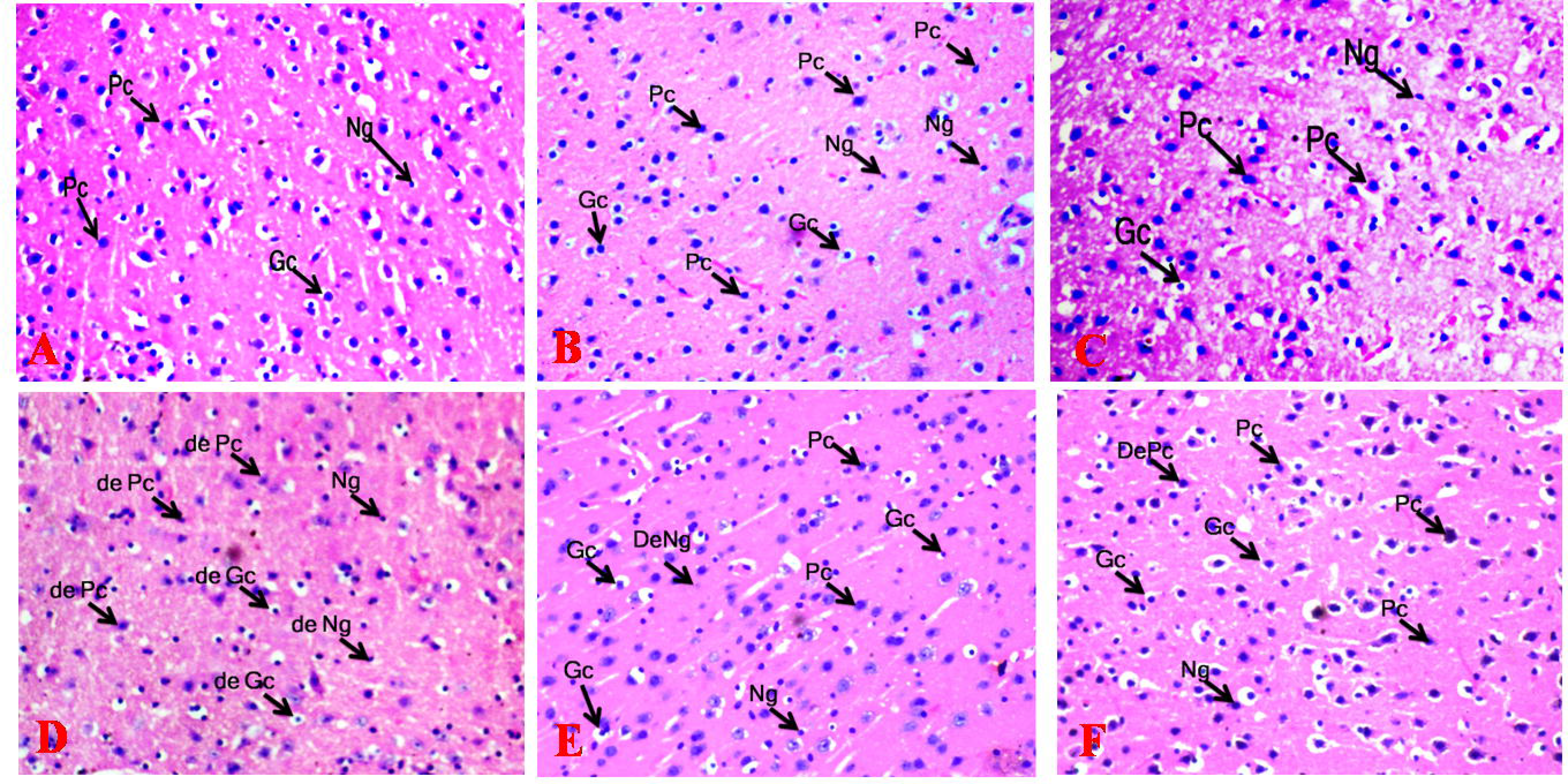
Effect of quercetin on cerebral morphology in cyclophosphamide (CYP)-administered rats. 9a) Control, 9b) Quercetin at 100 mg/kg body weight, 9c) Quercetin at 200 mg/kg body weight, 9d) Cyclophosphamide (CYP), 9e) CYP plus Quercetin at 100 mg/kg body weight, 9f) CYP plus Quercetin at 200 mg/kg body weight. Representative photomicrographs of haematoxylin and eosin (H&E) stained sections of the rat cerebral cortex show granule cells (Gc), pyramidal cells (Pc), neuroglia (Ng) and degenerating pyramidal cell (De Pc). H&E 160.

**Figure 10(a-f):**
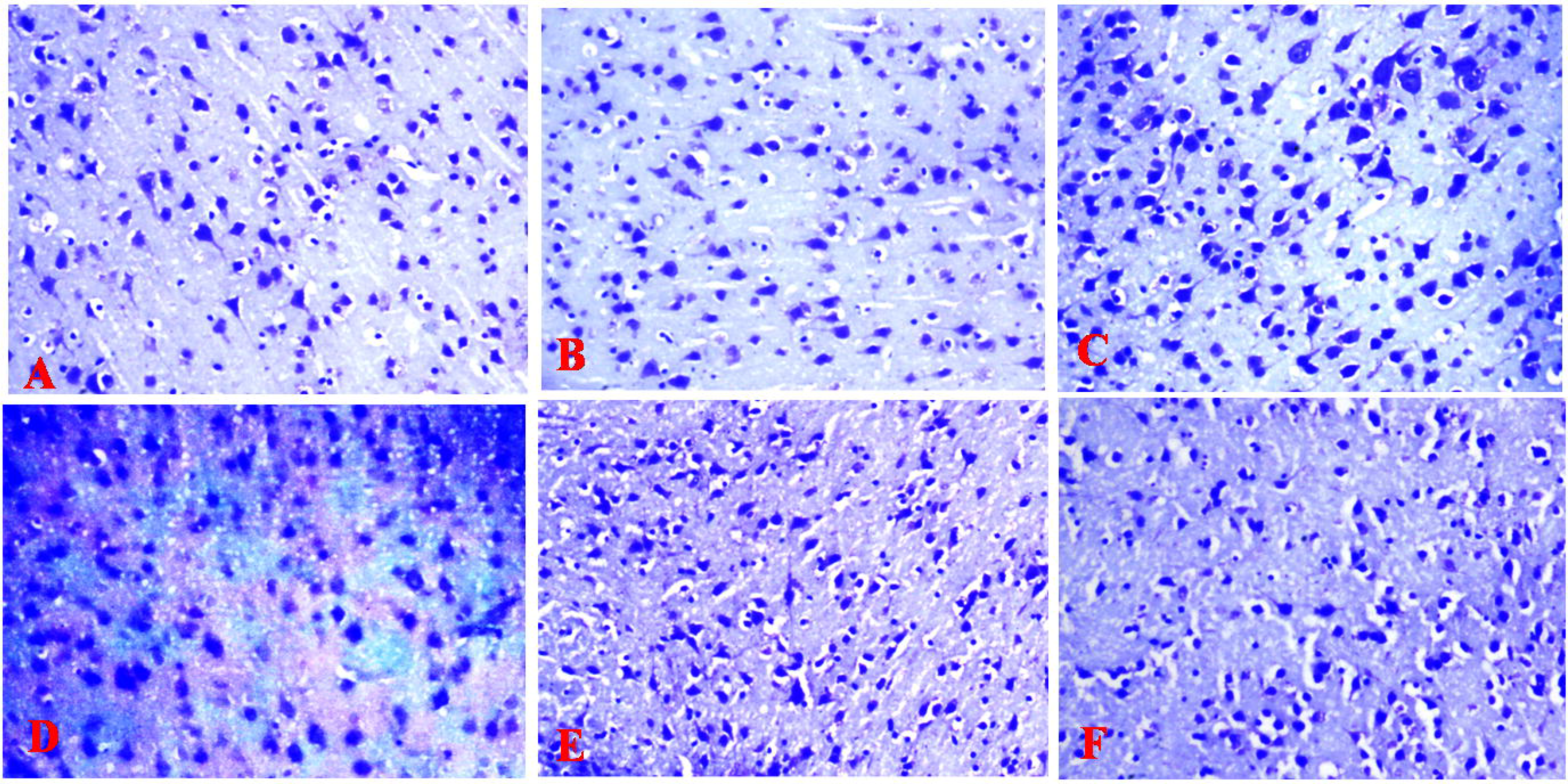
Effect of quercetin on cerebral morphology (cresyl fast violet stain) in cyclophosphamide (CYP)-administered rats. 10a) Control, 10b) Quercetin at 100 mg/kg body weight, 10c) Quercetin at 200 mg/kg body weight, 10d) Cyclophosphamide (CYP), 10e) CYP plus Quercetin at 100 mg/kg body weight, 10f) CYP plus Quercetin at 200 mg/kg body weight.. H&E 160.

### 3.9 Immunohistochemical staining of the Cerebrum

Immunohistochemical staining of the cerebral cortex (Fig. 11a-f) with anti- GFAP antibodies revealed the presence of few GFAP-positive immunoreactive astrocytes; although dispersed within the neuropil were brown cytoplasmic fibres scattered throughout the neuropil in rats in the control group (Fig. 11a). In groups administered quercetin (Figure 11b & 11c), a decrease in the density of immunoreactive astrocytes and their cytoplasmic fibres were observed. The group administered CYP (Figure 11 d was associated with a general decrease in neuronal cells; there was also a decrease in GFAP-reactive astrocytes, although these astrocytes showed some cellular hypertrophy, with thick cytoplasmic processes. However, individual astrocyte domains were fairly preserved without overlapping of astrocyte processes. In groups treated with quercetin reversal of the CYP induced changes were observed.

**Figure 11 (a–f):**
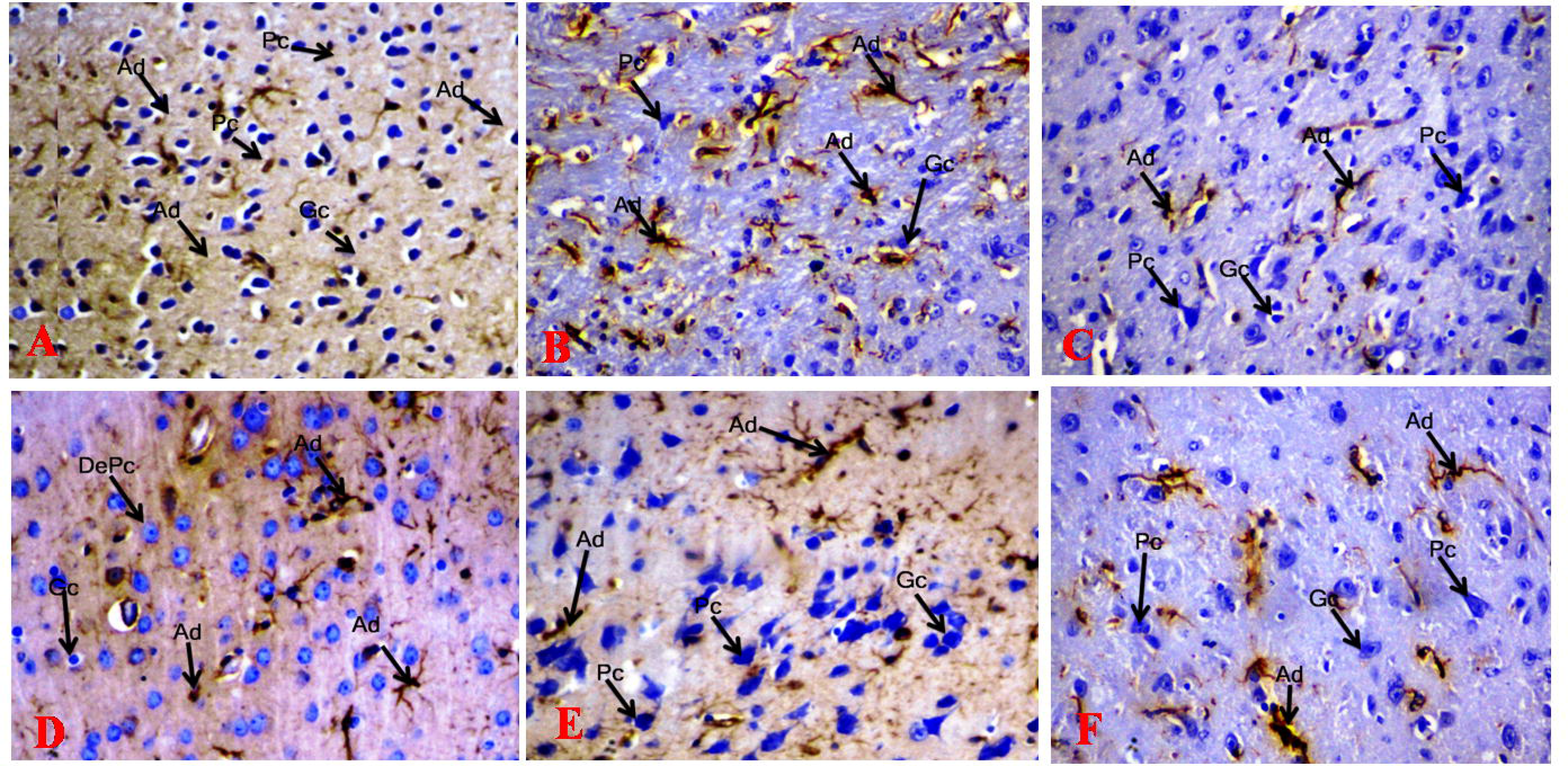
Effect of quercetin on glial fibrillary acidic protein (GFAP) immunohistochemistry of the cerebral cortex in cyclophosphamide (CYP)-administered rats.11a) Control, 11b) Quercetin at 100 mg/kg body weight, 11c) Quercetin at 200 mg/kg body weight, 11d) Cyclophosphamide (CYP), 11e) CYP plus Quercetin at 100 mg/kg body weight, 11f) CYP plus Quercetin at 200 mg/kg body weight. Representative photomicrographs of sections of the rat cerebral cortex following GFAP immunohistochemistry showing granule cells (Gc), pyramidal cells (Pc), and astrocytes dendrites (Ad). GFAP 160

## Discussion

This study investigated the possible protective effects of biflavonoid quercetin on changes in brain behaviour, biochemical and histomorphological markers of liver and kidney integrity, oxidant–antioxidant status, markers of inflammation, neurotransmitter levels and astrocyte immunoreactivity in healthy rats administered cyclophosphamide. The results showed that quercetin protected against CYP-induced changes in weight, food consumption, behavioural parameters, antioxidant status, inflammatory markers, brain neurotransmitters, and astrocytes immunoreactivity.

In this study, weekly body weight and percentage change in weight increased in the control group and decreased significantly in the groups administered cyclophosphamide. The effects on body weight observed with CYP in this study are consistent with the results of a number of other studies that had also reported that the administration of CYP caused significant weight loss (21, 43–45). On the other hand, treatment with quercetin mitigated CYP-induced weight loss Quercetin is a biologically active flavonoid that has been studied extensively for its effect on body weight (46–50). Results of these studies have varied significantly, with a few reporting quercetin’s ability to mitigate weight gain (46, 48), others observed no effect on body weight in normal sized animals (47, 49) in some other instances. In this study, groups administered quercetin alone showed no significant difference in body weight from control, supporting studies that had reported such effects. This would suggest that quercetin’s effect on weight only came into effect in situations of body weight abnormalities like disease related weight loss or conditions of obesity. In disease conditions as also observed in this study (with CYP related weight loss), quercetin’ s ability to reduce lipid peroxidation, increase antioxidant levels, and decrease inflammation even at the molecular level (51–53) are possible mechanisms through which it reversed CYP- induced weight loss. It could also reverse weight loss through its effects on food intake.

The administration of CYP in this study was associated with a significant reduction in food intake while treatment with quercetin reversed CYP induced decrease in food intake. In groups administered quercetin alone, no significant difference in food intake was observed, supporting our earlier observations that quercetin’s effect on the body was to restore normalcy. A few studies have also reported that quercetin did not increase energy consumption (54). In the CYP groups, quercetin supplementation reversed CYP induced decrease in food intake, consistent with the results of other studies that had observed quercetin’s ability to reverse food intake in a disease model (55). Quercetin’s effects on body weight and food intake can also be attributed to its ability to influence central processes that regulate energy homeostasis (55).

In vertebrates, the liver is a very important regulator of energy homeostasis, this it achieves through its ability to sense nutrient availability and alter energy production. The beneficial effect of quercetin in mitigating liver injury has been reported severally (56–58), with a few of these studies attributing its hepatoprotective effects to its ability to increase the activities of metabolising enzymes, antioxidant status and decreasing lipid peroxidation and the expressions of proinflammatory cytokines (56, 57). In this study the administration of quercetin alone did not alter the activities of hepatic enzymes although it increased total antioxidant capacity however, in groups administered CYP there was an increase in the activities of AST and ALT a decrease in total antioxidant capacity and an increase in lipid peroxidation. The effects of CYP on serum levels of hepatic enzymes, antioxidant capacity and lipid peroxidation are consistent with the results of previous studies (59–62). Injury to the cellular components of the liver has been reported to play important roles in hepatic cell death (63). Consequently level of hepatic enzymes like AST and ALT increase due to their release, therefore high serum levels of these enzymes are generally indicative of liver injury. This notion is supported by histological evidence of hepatic injury in this study. The administration of quercetin in the CYP group of animals was associated with a reversal of the biochemical and histological effects induced by CYP. The beneficial effects of quercetin in protecting against liver injury have been attributed to its ability to enhance antioxidant capacity and decrease pro-oxidant effects (63). There have also been reports that quercetin can scavenge superoxide, hydroxyl, peroxyl and alkoxyl radicals in tissues (57, 63).

Studies have continued to report the benefits of flavonoids and flavonoid-rich compounds in modulating inflammatory activity and in protecting against organ injury (64–66). The beneficial effects of quercetin could also be attributed to its effects on other organs like the kidney that are also important in regulating energy metabolism. The proinflammatory and nephrotoxic effects of CYP have also been reported (67–69). In consistency with these reports, this study observed derangement of urea and creatinine levels and an increase in proinflammatory cytokines (TNF-α and IL 1β); while levels of antiinflammatory cytokines (IL-10) decreased. Also, in this study, the administration of CYP was associated with histological evidence of renal injury. Dietary supplementation with quercetin was associated with reversal of urea and creatinine derangements as well as a decrease in the levels of proinflammatory and an increase in the levels of anti-inflammatory cytokines. The nephroprotective effects of quercetin have been reported (70 71). The nephroprotective effects of quercetin have been linked to its ability to inhibit the formation of malondialdehyde, scavenge reactive oxygen species, and activate the expression of nuclear factor erythroid 2-related factor 2 and renal heme oxygenase 1, which are important in boosting the production of antioxidant moieties (72).

Also in this study, the protective effect of quercetin on neurobehaviour and brain histomorphology was observed. Neurobehavioural paradigms are non-invasive models that are employed for the assessment of normal central nervous system function or to investigate the effects of drug and/or drug candidates on the functioning of the central nervous system in health or disease [73]. Several studies have demonstrated that cyclophosphamide, a commonly used anti cancer chemotherapeutic agent has neurotoxic effects (74–80). In this study, we observed that the administration of cyclophosphamide was associated with central excitation, increased anxiety and memory impairment. Neurochemicals such as acetylcholine, dopamine, and brain derived neurotropic factor decreased significantly, while serotonin levels increased. Cyclophosphamide administration was also associated with morphological evidence of neuronal injury. Overall, the administration of cyclophosphamide has neurotoxic effect. Reports from a study by Flanigan et al (81) showed no significant effect of CYP on locomotor activity and motor coordination; although an increase in ambulatory behaviour was observed during assessment of mouse behaviour in their home cages. The effects observed on locomotor activity by Flanigan et al (81) contrast the effects observed in this study. However, more consistently reported are the memory impairing effects of CYP (75, 78, 81) which was also observed in this study. There have been reports that the ability of cyclophosphamide to cross the blood brain barrier and induce oxidative stress is one of the many ways through which it induces neurotoxicity (78). Current research is geared towards evaluating the possible beneficial effects of antioxidant compounds like quercetin on mitigating the neurotoxic effects of CYP. Also in this study, cyclophosphamide was associated with neuronal injury and a decrease in immunoreactive astrocytes. This is consistent with the results of Ibrahim et al (82), which reported evidence of neuronal injury in the hippocampus following cyclophosphamide administration. However, results of this study differed in that they observed an increase in immunoreactive astrocytes compared to a decrease observed in this study. The 2001 study by Shanani et al (83) observed that cyclophosphamide administration was associated with a decrease in immunoreactive astrocytes in a rat model of amyotrophic lateral sclerosis. In our study, while there was a decrease in immunoreactive astrocytes in the cerebral cortex, the cytoplasmic processes of these astrocytes were thicker than observed in the control group suggesting the possibility of an increase in activity of the few astrocytes observed.

Also, in this study, quercetin supplementation concentration-dependently increased horizontal locomotion, self grooming and radial arm maze spatial working memory, while at both concentrations of quercetin, an anxiolytic effect and increase in Y maze spatial working memory was observed. Quercetin’s ability to reverse brain injury has been reported severally (85–90). While most studies have examined its benefits in Alzheimer’s disease, only a few studies have examined its potential benefits in chemotherapy induced neurotoxicity (91). In this study, we observed that quercetin reversed CYP induced changes in behaviour and brain injury through its ability to modulate neurotransmitter levels and glia fibrillary acidic protein immunoreactivity in the cerebral cortex and hippocampus. Quercetin’s antioxidant effects have also been postulated as a potential mechanism for its effects on the brain. Although we did not examine the effects of quercetin supplementation on brain levels of inflammatory and oxidative stress markers, the examination of its peripheral effect revealed a reversal of CYP-induced changes in serum inflammatory markers. Studies examining the mechanisms responsible for quercetin’s effect on the brain have also listed quercetin’s ability to rescue apoptotic pathways and increase neurogenesis as potential mechanisms. In this study, reversal of morphological changes observed due to CYP could be due to reduction in the destructive activity on astrocytes.

## Conclusion

In conclusion, the results of the current study highlight a possible application of quercetin as an adjunct in cancer chemotherapy, especially for the purpose of mitigating unwanted effects of chemotherapy. However, one area that is also likely to need further clarification is that of its possible direct interaction with the chemotherapy agents. This further research will likely point us in the direction that we are likely to go, while we wait to realise the benefits that may accrue from the use of quercetin in this context.

## Author Contribution Statement

A.Y.O and O.J.O conceived and designed research. AYO, FOO and OJO conducted experiments and analyzed data. A.Y.O and O.J.O wrote the manuscript. All authors read and approved the manuscript.

## Ethics Approval and Consent to Participate

All procedures performed on the animals were in accordance with approved protocols of the Faculty of Basic Medical Sciences, Ladoke Akintola University of Technology, Ogbomoso, Oyo State, Nigeria and within the provisions for animal care and use as prescribed by the scientific procedures on living animals, European Council Directive (EU2010/63).

## Human and Animal Rights

No humans were used in this study. All procedures performed on the animals were in accordance with approved protocols of the Ladoke Akintola University of Technology

## Consent for Publication

Not Applicable.

## Availability of Data and Materials

The datasets generated during and/or analysed during the current study are available from the corresponding author on reasonable request.

## Conflict of interest

All authors of this paper declare that there is no conflict of interest related to the content of this manuscript.

## Source of funding

This research did not receive any specific grant from agencies in the public, commercial, or not-for-profit sectors.

## Notes

### Competing Interest Statement

The authors have declared no competing interest.

